# Large-scale analysis of genome and transcriptome alterations in multiple tumors unveils novel cancer-relevant splicing networks

**DOI:** 10.1101/023010

**Authors:** Endre Sebestyén, Babita Singh, Belén Miñana, Amadís Pagès, Francesca Mateo, Miguel Angel Pujana, Juan Valcárcel, Eduardo Eyras

## Abstract

Alternative splicing is regulated by multiple RNA-binding proteins and influences the expression of most eukaryotic genes. However, the role of this process in human disease, and particularly in cancer, is only starting to be unveiled. We systematically analyzed mutation, copy number and gene expression patterns of 1348 RNA-binding protein (RBP) genes in 11 solid tumor types, together with alternative splicing changes in these tumors and the enrichment of binding motifs in the alternatively spliced sequences. Our comprehensive study reveals widespread alterations in the expression of RBP genes, as well as novel mutations and copy number variations in association with multiple alternative splicing changes in cancer drivers and oncogenic pathways. Remarkably, the altered splicing patterns in several tumor types recapitulate those of undifferentiated cells. These patterns are predicted to be mainly controlled by *MBNL1* and involve multiple cancer drivers, including the mitotic gene *NUMA1*. We show that *NUMA1* alternative splicing induces enhanced cell proliferation and centrosome amplification in non-tumorigenic mammary epithelial cells. Our study uncovers novel splicing networks that potentially contribute to cancer development and progression.

## Introduction

Alternative splicing alterations are emerging as important signatures to further understand tumor formation and to develop new therapeutic strategies (Grosso and Carmo-Fonseca 2014). Specific alternative splicing changes that confer tumor cells with a selective advantage may be caused by mutations in splicing regulatory sequences (Dorman et al. 2014) and/or regulatory factors (Brooks et al. 2014). Various splicing factors have been described to be mutated in cancer, including *SF3B1, SRSF2, ZRSR2, U2AF1* in myelodysplastic syndromes and lymphoid leukemias (Yoshida et al. 2011), *RBM10* and *U2AF1* in lung tumors (Brooks et al. 2014) (Imielinski et al. 2012) and *SF3B1* in breast tumors (Maguire et al. 2015). These mutations generally impair the recognition of regulatory sites, thereby affecting the splicing of multiple genes, including oncogenes and tumor suppressors (Kim et al. 2015). On the other hand, increasing evidence shows that changes in the relative concentration of splicing factors can also trigger oncogenic processes. For instance, splicing factors from the SR and hnRNP families are overexpressed in multiple tumor types and induce splicing changes that contribute to cell proliferation (Karni et al. 2007) (Golan-Gerstl et al. 2011). Similarly, downregulation of splicing factors that act as tumors suppressors has also been observed (Wang et al. 2014) (Zong et al. 2014).

Importantly, specific alternative splicing events can substantially recapitulate cancer-associated phenotypes linked to mutations or expression alterations of splicing factors. This is the case *of NUMB*, for which the reversal of the splicing change induced by *RBM10* mutations in lung cancer cells can revert the proliferative phenotype (Bechara et al. 2013). Events that contribute to cancer are often controlled by multiple factors, like the exon skipping event *of MST1R* involved in cell invasion, which is controlled by *SRSF1* (Ghigna et al. 2005), *HNRNPA2B1* (Golan-Gerstl et al. 2011), *HNRNPH1* and *SRSF2* (Moon et al. 2014). Furthermore, some events may be affected by both mutations and expression changes in splicing factors. For instance, mutations in *RBM10* or downregulation of *QKI* lead to the same splicing change in *NUMB* that promotes cell proliferation (Bechara et al. 2013) (Zong et al. 2014). Alternative splicing changes that characterize and contribute to the pathophysiology of cancer (Sebestyén et al. 2015) are thus potentially triggered by alterations in a complex network of RNA binding proteins, which remains to be comprehensively described. To elucidate the complete set of alterations in these factors and how they globally affect alternative splicing that may contribute to cancer, we analyzed RNA and DNA sequencing data from The Cancer Genome Atlas (TCGA) project for 11 solid tumor types.

## Results

### *RBPs* are frequently deregulated and characterize tumor types

Using TCGA data for 11 solid tumor types (Supplemental Table S1), we analyzed the differential gene expression between normal and tumor sample pairs of 1348 genes encoding known and predicted RNA binding proteins (RBPs) (Supplemental Table S2) (Methods). The majority of these genes (1143, 84,8%) show significant differential expression in at least one tumor type (Supplemental Fig. S1) (Supplemental Table S3). Examining in detail 162 RBP genes annotated as known or putative splicing factors (SFs), they can be separated into three groups. One group is frequently upregulated, another one downregulated, and a third one shows opposite patterns in the three kidney tumor types (KICH, KIRC, KIRP) compared to other tumor types (Fig. 1A). Moreover, 132 (80%) of them are differentially expressed in at least one tumor type and 45 were previously associated with oncogenic or tumor suppressor activities (Fig. 1A, labeled in red) (Supplemental Table S4). We also found apparent discrepancies with previous literature. For instance, although *SRSF5* was described as oncogenic (Huang et al. 2007) it is downregulated in six tumor types; and *TRA2B*, reported as oncogenic in breast cancer (Watermann et al. 2006), is upregulated in LUSC but downregulated in KICH and thyroid carcinoma (THCA). Also, the oncogenic *SRSF2, SRSF3* and *SRSF6* (Xiao et al. 2007; Jia et al. 2010; Jensen et al. 2014) are downregulated in KICH, while the oncogenic *SRSF1* and the tumor suppressor *RBM4* do not show any significant expression changes. New patterns also emerge, including upregulation of genes from the RBM family and downregulation of the genes from the *MBNL* family (Fig. 1A). Unsupervised clustering of all the 4442 tumor samples using normalized expression values per sample for SFs (Supplemental Fig. S2) or for all RBPs (Supplemental Fig. S3) largely separates samples by tumor type. This pattern is reproduced when using a different gene set of similar size (Supplemental Fig. S3). Closer inspection of the BRCA and COAD samples reveals that the expression patterns of RBPs also reproduce tumor subtypes (Supplemental Figs. 4 and 5). Collectively, expression analyses indicate that RBPs are frequently and specifically altered in human tumors and they represent one of the multiple expression profiles that characterize the intrinsic properties of the tumor types and their tissue of origin.

**Figure 1.**
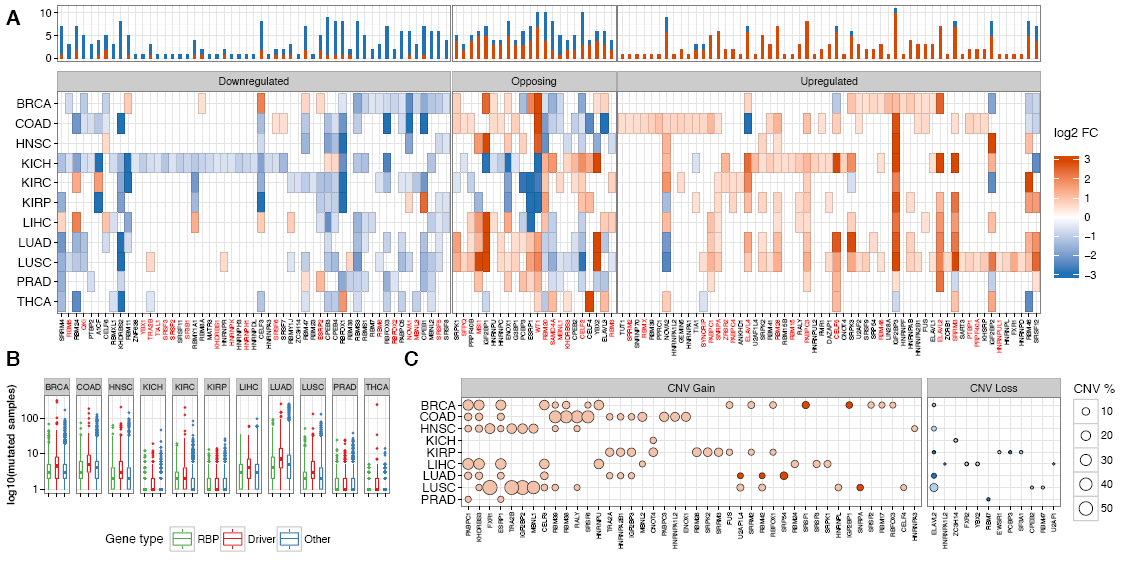
Cancer alterations in splicing factors. **(A)** Up- (red) and downregulation (blue) patterns splicing factors (x-axis) in the different tumor types (y-axis) compared to normal samples. Only the 132 SFs (out of 162 tested) with differential expression are shown. The color intensity indicates the log2-fold change (log2 FC). The bar plot above indicates the frequency of tumor types with up- (red) or down- (blue) regulation for each factor. Factors are clustered into three groups according to whether they show frequent downregulation (Downregulated) or upregulation (Upregulated) in tumors, or whether they tend to show an opposite pattern between the three kidney tumors (KICH, KIRC, KIRP) and the rest of tumor types (Opposing). Factors previously described to have oncogenic or tumor-suppressing activities (Supplemental Table S3) are labeled in red. *SF3B1, SRPK1, SRPK2* and *SRPK3* were included for comparison. **(B)** Number of samples in log10 scale (y axis) in which RBPs (green), driver genes (red) (Supplemental Tables S5 and S6) and the rest of genes (blue) show mutations in each tumor type (x axis). Distributions are represented as box plots, with outliers represented as dots. All comparisons of drivers vs. RBPs or drivers vs. others are significant (one-sided Wilcoxon test p-values < 1.7e-05). Comparisons of others vs. RBPs are significant (one-sided Wilcoxon test p-values < 0.05), except for LUAD, LUSC and PRAD. Drivers were extracted from the literature (Methods). **(C)** Copy number variation (CNV) gains (left panel) and losses (right panel) of the tested splicing factors. The size of the circle corresponds to the proportion of samples with CNVs and darker colors represent cases where more than 50% of the CNVs are focal. Only those CNVs with a frequency of amplification > 5% or deletion > 1% are shown.

### *RBPs* deregulation is partially driven by genomic alterations

To further define the extent to which RBP genes are altered in human cancer, the TCGA data was analyzed for protein-affecting mutations and copy number variations (CNVs). Although most of the RBP genes are mutated in tumors (Supplemental Table S4), they are generally mutated in fewer samples compared to candidate cancer drivers (listed in Supplemental Tables S5 and S6) and other genes (Fig. 1B). We confirmed 5,4% (25/458) of LUAD tumor samples to have protein-affecting mutations for *RBM10*, in agreement with previous studies (Imielinski et al. 2012). Using this case as reference, we observed 205 (15.2%) of all RBPs (13 or 8% of the 162 SFs) mutated in more than 5% of samples in a given tumor type (Table 1) (Supplemental Table S3). In general, there is a weak association between mutations and expression changes, whereas a number of genes show mutual exclusion of mutations and expression changes (Table 1) (Supplemental Table S3). On the other hand, CNVs are highly recurrent across samples, with gains more frequent than losses (Fig. 1C) (complete list of CNVs available in Supplemental Table S3). Upregulated RBPs had generally more CNV gains than non-regulated RBPs (Mann-Whitney test p-value < 2.2e-16) and 90% of the upregulated RBPs had CNV gains in more than 50% of the samples from the same tumor type. The splicing factors *ESRP1, PABPC1* and *KHDRBS3*, which are near the reported 8q24 amplification (The Cancer Genome Atlas Network 2012b), show CNVs in BRCA, COAD, HNSC, LIHC and LUAD (Fig. 1C), with *ESRP1* and *PABPC1* showing frequent association with upregulation (Table 1). We also observed the frequent amplification of *TRA2B, IGF2BP2* and *FXR1* in HNSC and LUSC, as reported before (The Cancer Genome Atlas Network 2012a, 2015). However, the association with upregulation is only observed in LUSC (Table 1). CNV losses are less frequent than gains and show weaker associations with downregulation (Table 1) (Supplemental. Table S3). Finally, only a low fraction of the CNVs were detected as focal (Table 1) (Supplemental Table S3). These results indicate that many of the detected expression alterations of RBPs may be explained by CNVs, although most often in association with large chromosomal events, thereby highlighting the potential relevance of these alterations in shaping the tumor phenotypes.

**Table 1.**
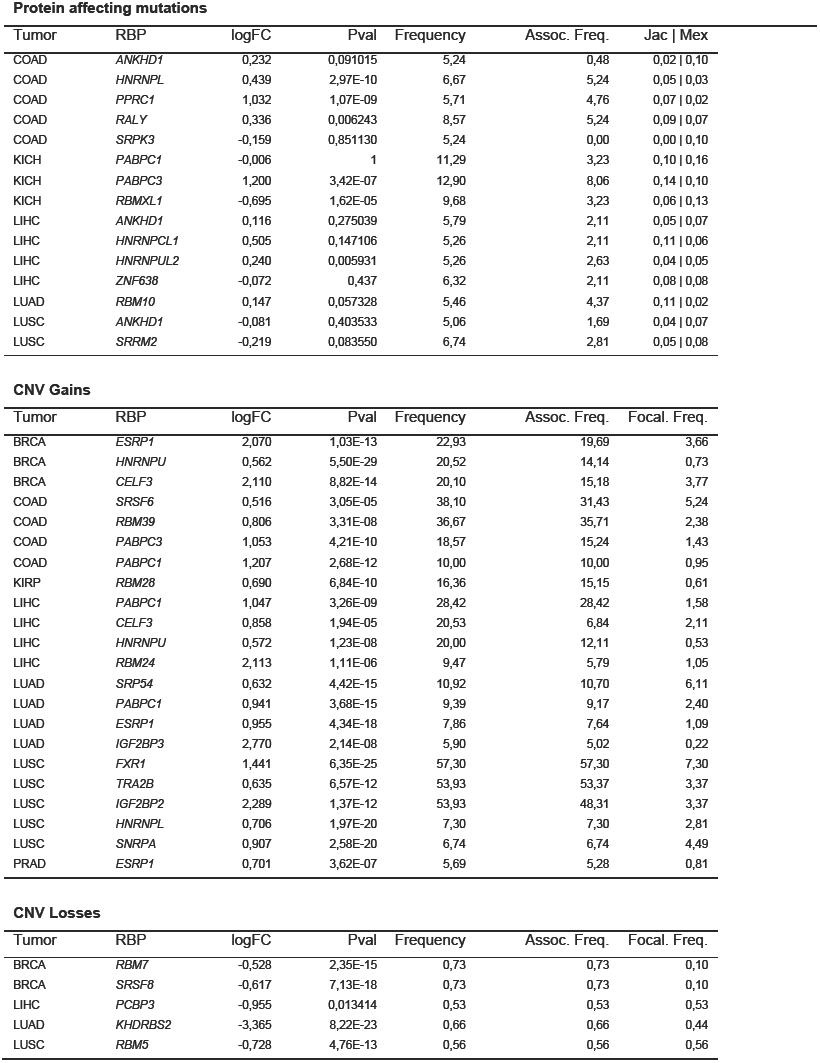
Association of mutations and CNVs with expression changes. For each RBP and each tumor type we indicate the log2-fold change (logFC) and adjusted p-value (Pval) of the differential expression analysis between tumor and normal samples, the frequency of the alteration (Frequency) and the association of the alteration with the expression alteration (Assoc. Freq.). For mutations we show those cases with mutation frequency >5% and with a Jaccard (Jac) or mutual exclusion (Mex) score > 0.05. For CNV gains we show those cases that have significant upregulation, CNV gain and association frequencies in > 5% of the samples, with one or more of the CNVs being focal (Focal. Freq). For CNV losses we show those with expression frequency and association with down regulation in > 0.5% of the samples, with one or more of the CNVs being focal (Focal. Freq). Complete data for all RBPs is given in the Supplemental Table S3.

### *RBP* mutations associate with abnormal splicing patterns in cancer

Next, we investigated the patterns of differential splicing that may occur as a consequence of the alterations described above. To determine those possibly related to expression alterations in RBPs, we first evaluated the significant splicing changes in tumors compared to normal tissue, considering five major event types: skipping exon (SE), alternative 5’ splice-site (A5), alternative 3’ splice-site (A3), mutually exclusive exon (MX) and retained intron (RI) events (Fig. 2A) (Supplemental Tables S7 and S8). We examined the splicing patterns of cancer drivers in more detail. From the 937 drivers collected (Supplemental Tables S5 and S6), 653 (69.7%) have annotated events and 292 (31.2%) have at least one differentially spliced event, with a number of them occurring in multiple tumor types (Supplemental Fig. S6). Moreover, 7 of the 11 tumor types show enrichment of differentially spliced events in drivers (Fig. 2B, in red) (Supplemental Table S9). Additionally, various cancer hallmarks (Liberzon et al. 2015) are enriched in differentially spliced events but not in differentially expressed genes (Fig. 2C) (Supplemental Fig. S7) (Supplemental Table S10). These results suggest that alternative splicing contributes to cancer development independently of mutations or expression alterations.

**Figure 2.**
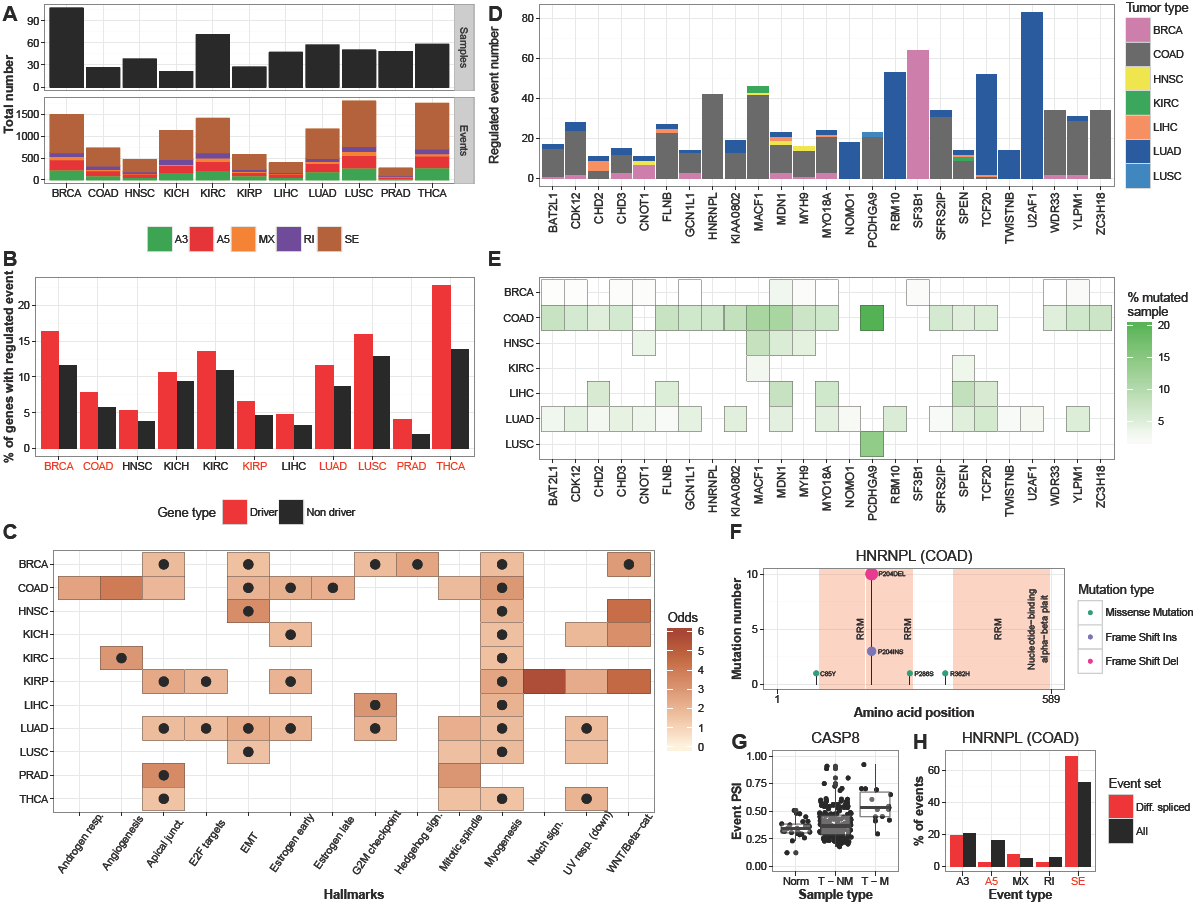
Differentially spliced events in tumors. **(A)** Upper panel: number of paired-samples used per tumor type. Lower panel: number of differentially spliced events per tumor type compared to normal samples, split according to type of event: alternative 3’ splice-site (A3), alternative 5’ splice-site (A5), mutually exclusive exon (MX), retained intron (RI) and skipping exon (SE) (Supplemental Table S6). **(B)** Proportion of driver and non-driver genes with differentially spliced events. We indicated in red those tumors for which the enrichment is significant. **(C)** Cancer hallmarks (x-axis) that are enriched (Fisher test p-value < 0.05) in differentially spliced events in each tumor type (y-axis). The color indicates the odds ratio of the enrichment. Hallmarks that are also enriched according to gene expression are indicated with a black dot. **(D)** Number of differentially spliced events related to protein-affecting mutations in RBP genes color-labeled by tumor type. Only cases with at least 10 associated differentially spliced events are shown. *SF3B1* is included for comparison. **(E)** Proportion of samples per tumor type with protein-affecting mutations in RBP genes with at least 10 associated differentially spliced events. **(F)** Number of protein-affecting mutations (y-axis) along the HNRNPL protein (x-axis), color-labeled according to whether they are substitutions, insertions or deletions. Protein domains are indicated in light red. **(G)** Distribution of PSI values for the A5 event in *CASP8* associated to the mutations of *HNRNPL* in COAD, separated into normal samples (Norm), tumor samples without protein-affecting mutations (T – NM), and tumor samples with protein-affecting mutations (T – M). **(H)** Enrichment or depletion of specific event types in association to mutations in *HNRNPL* (red bars) compared to the overall proportions of events (black bars). Significant differences (p < 0.05, Fisher test) are labeled in red. Contingency tables are provided in Supplemental Table S12.

To determine the splicing changes related to mutations in RBPs, we compared the inclusion levels (percent spliced in, PSI) of the events between samples with or without protein-affecting mutations for each RBP (Figs. 2D and 2E) (Supplemental Fig. S8) (Supplemental Table S11). We found less differentially spliced events compared with the tumor-normal comparison and some of the detected events affect cancer drivers (Table 2). *HNRNPL* had 16 mutations in COAD, 13 of them indels in an RNA recognition motif; causing frameshifts (Fig. 2F). These mutations had 42 events associated with them (Z-score = 41.35), including the cancer driver *CASP8* (Fig. 2G). In COAD we also found the putative RBP *MACF1* (Baltz et al. 2012) with mutations along the entire protein (42 events, Z-score = 45.35) (Supplemental Fig. S9). *MACF1* is a component of the WNT-pathway known to affect splicing (Bordonaro 2013). In LUAD, we found 53 events associated to *RBM10* (Z-score = 52.35), 83 events associated to *U2AF1* (Z-score = 81.35), and 49 events associated to the transcription factor and putative RBP *TCF20* (Castello et al. 2012) (Z-score = 50.35). Most of the *TCF20* alterations consist of an insertion in a Glycine-rich region at the N-terminus (Supplemental Fig. S9). As a comparison, we analyzed *SF3B1* and found 64 differentially spliced events (Z-score = 63.35) in BRCA, despite being mutated in only 1.7% of the tested samples (16 out of 956 for which we had mutation and RNA-Seq data) (Supplemental Fig. S9). Notably, compared with the proportions of the different event types, there is a significant enrichment of A3 events associated with *SF3B1* (Fisher’s test p-value = 1.45E-11), of A5 events for *TCF20* (p-value = 2.09E-6), and of SE events for *RBM10* (p-value = 0.003) (Supplemental Fig. S9) (Supplemental Table S12). There is also a significant depletion of SE events for *TCF20* (p-value = 0.0014), of SE events for *SF3B1* (p-value = 2.24E-8), and of A5 events for *U2AF1* (p-value = 1.67E-3) and *HNRNPL* (p-value = 0.005) (Fig. 2H). Furthermore, we confirmed 17 (20%) of the previously detected differentially spliced genes for *SF3B1* (Maguire et al. 2015), 32 (38%) for *U2AF1* (Brooks et al. 2014) and 21 (30%) for *RBM10* (Brooks et al. 2014) (Supplemental Table S13). In summary, we validated known RBP mutations affecting alternative splicing and described new ones, whose mutations potentially affect splicing patterns. Our analyses reveal a rich source of new information about alternative splicing events associated to RBP alterations with a potential relevance in cancer.

**Table 2.** Events in cancer drivers associated to protein-affecting mutations in RBPs. For each tumor type and for each RBP with more than 10 associated differentially spliced events, we indicate the events in cancer drivers predicted to have a significant splicing change. The table includes the PSI change between mutated and non-mutated samples (ΔPSI) and the p-value of the comparison after correcting for multiple testing (Pval). *SF3B1* is included for comparison. Coordinates for the events are given in Supplemental Table S7.

| Tumor type | Gene | Cancer driver | Event type | ΔPSI | Pval |
| --- | --- | --- | --- | --- | --- |
| BRCA | *CDK12* | *TFEB* | SE | -1,00 | 0,0001 |
| BRCA | *MDN1* | *EBF1* | SE | -0,23 | 0,0343 |
| BRCA | *SF3B1* | *BCL2L1* | A3 | 0,23 | 4,14E-06 |
| BRCA | *SF3B1* | *MEF2A* | SE | -0,21 | 0,0026 |
| COAD | *BAT2L1* | *MLH1* | SE | 0,70 | 0,0109 |
| COAD | *CHD2* | *CAST* | SE | -0,15 | 0,0396 |
| COAD | *HNRNPL* | *CASP8* | A5 | 0,17 | 0,0271 |
| COAD | *KIAA0802* | *MKNK2* | A3 | -0,13 | 0,0340 |
| COAD | *KIAA0802* | *MYH11* | MX | -0,14 | 0,0004 |
| COAD | *MACF1* | *TJP2* | SE | 0,16 | 0,0251 |
| COAD | *MYH9* | *AURKA* | A5 | 0,16 | 0,0462 |
| COAD | *MYO18A* | *MDM4* | A3 | 0,54 | 0,0462 |
| COAD | *MYO18A* | *PDCD1LG2* | A5 | 0,21 | 0,0403 |
| COAD | *SPEN* | *TTLL9* | SE | -0,17 | 0,0118 |
| COAD | *YLPM1* | *CD44* | SE | 0,12 | 0,0268 |
| COAD | *ZC3H18* | *ACSL6* | MX | -0,84 | 0,0114 |
| COAD | *ZC3H18* | *MBD1* | SE | -0,11 | 0,0173 |
| HNSC | *DSP* | *PAX5* | MX | -0,12 | 0,0034 |
| HNSC | *EPPK1* | *TAF1* | SE | 0,24 | 0,0007 |
| HNSC | *MYH9* | *PTCH1* | SE | -1,00 | 0,0058 |
| LIHC | *CHD2* | *BUB1B* | SE | 1,00 | 0,0020 |
| LIHC | *EPPK1* | *NEDD4L* | SE | -0,26 | 0,0339 |
| LIHC | *HUWE1* | *HLA-A* | A5 | 0,35 | 0,0281 |
| LUAD | *CHD2* | *SELP* | A3 | -1,00 | 0,0008 |
| LUAD | *NOMO1* | *TFG* | A5 | -0,23 | 0,0238 |
| LUAD | *RBM10* | *BLM* | SE | 0,13 | 0,0340 |
| LUAD | *RBM10* | *CTNND1* | A3 | -0,34 | 0,0064 |
| LUAD | *RBM10* | *MUC1* | SE | 0,13 | 0,0333 |
| LUAD | *RBM10* | *WNK1* | A3 | 0,22 | 0,0033 |
| LUAD | *TWISTNB* | *CLIP1* | MX | -0,20 | 0,0435 |
| LUAD | *U2AF1* | *BCOR* | SE | -0,12 | 0,0351 |
| LUAD | *U2AF1* | *CHCHD7* | A3 | -0,11 | 0,0005 |
| LUAD | *U2AF1* | *CTNNB1* | A3 | 0,26 | 0,0002 |
| LUAD | *U2AF1* | *CTNNB1* | RI | 0,19 | 0,0004 |
| LUAD | *U2AF1* | *MUC1* | MX | -0,26 | 0,0472 |
| LUAD | *U2AF1* | *PATZ1* | A3 | -0,15 | 0,0347 |
| LUAD | *U2AF1* | *PCM1* | SE | 0,12 | 0,0160 |
| LUAD | *U2AF1* | *RIPK2* | SE | -0,26 | 0,0008 |
| LUAD | *U2AF1* | *RIT1* | SE | -0,13 | 0,0018 |

### Common and specific cancer patterns of differential splicing mediated by RBPs

The above results suggest that mutations in RBPs are not the single cause of splicing changes in tumors. Therefore, we further characterized the splicing changes between tumor and normal samples to determine their association with the expression of RBPs. We first identified common patterns of splicing changes between pairs of tumor types by selecting events with a strong correlation with a differentially expressed SF in the two tumor types (Methods). PSI changes (ΔPSI) for these events show high correlation between tumor pairs (Fig. 3A) and indicate potential common regulators (Fig. 3B) (Supplemental Fig. S10) (Supplemental Table S14). BRCA and LUAD share 229 events associated with various factors, including *QKI* and *SRSF5* (Fig. 3B, upper left panel), whereas KIRC and PRAD share 78 anti-correlating events associated with *ESRP2* and *MBNL1* (Fig. 3B, upper right panel). *RBM47*, described as tumor suppressor (Vanharanta et al. 2014), appears as the main common SF between KIRC and HNSC with 61 associated events (Fig. 3B, lower left panel). HNSC and LUSC share 141 events, 63 of them associated with *RBM28* (Fig. 3B, lower right panel). These results suggest functional connections between RBPs and the splicing changes detected in cancer. Next, we also studied whether there are tumor specific events (Methods) and found 380 events that largely separate the 4442 tumor samples by type (Fig. 3C) (Supplemental Fig. S11) (Supplemental Table S15). These splicing changes may be indicative of tumor-type specific oncogenic mechanisms.

**Figure 3.**
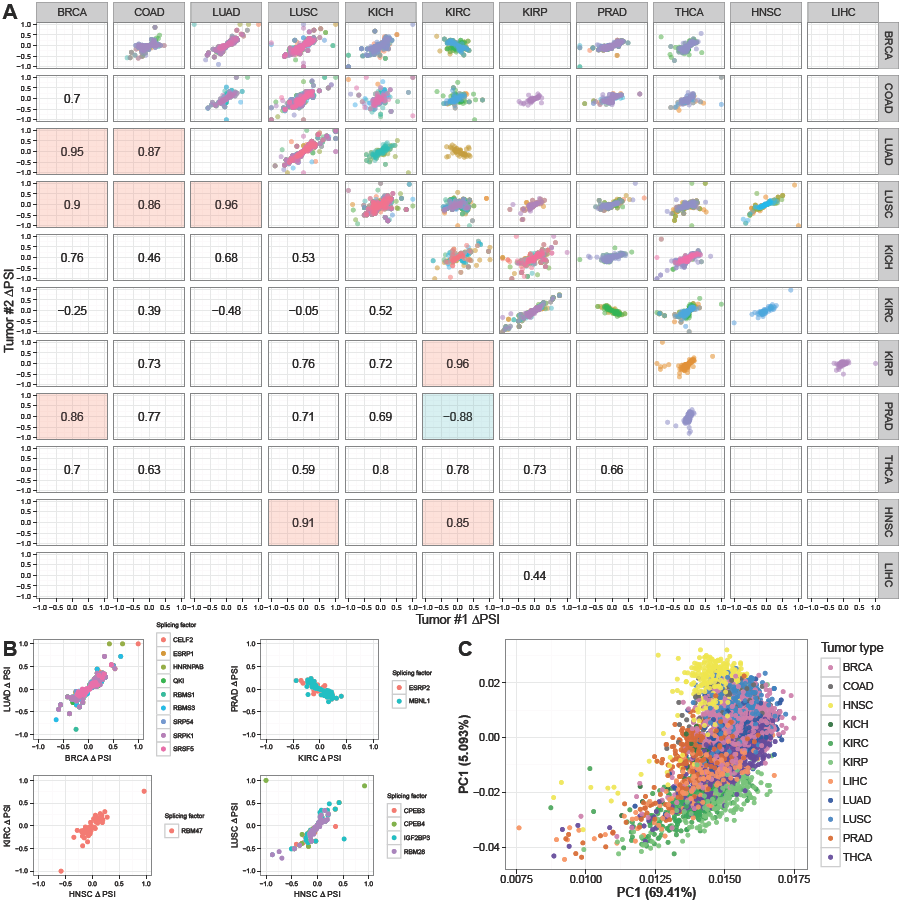
Common and specific events in tumors. **(A)** Common events and splicing factors between pairs of tumor types. For each pair of tumor types and for each splicing factor differentially expressed in both tumor types, we plot the correlation of ΔPSI values for events that have a correlation of |R| > 0.5 (Spearman) with these splicing factors in both tumor types. Only factors with more than 50 associated events in both tumor types are shown. Each event is only plotted once and the color of the plot corresponds to the most common correlating splicing factor. Correlations between ΔPSI values are indicated. In red or green, we highlight those higher than 0.8 or lower than −0.8, respectively. **(B)** ΔPSI correlations for the pairs LUAD – BRCA, PRAD – KIRC, KIRC – HNSC, and LUSC – HNSC, for the common events separated according to their potential splicing factor regulators. Events associated to more than one factor are represented with jitter. **(C)** Principal Component Analysis (PCA) plot of 380 tumor specific alternative splicing events colored by tumor type.

### Enriched RBP motifs in differentially spliced events of cancer

To further understand the link between the observed splicing patterns and the RBPs, we tested the enrichment of binding motifs in differentially spliced events in each tumor type. We assigned binding motifs from RNAcompete (Ray et al. 2013) to 104 of the analyzed RBPs and tested their enrichment (see Methods and Supplemental Figs. S12 and S13). Considering enriched motifs of differentially expressed RBPs in the same tumor type, we observed that motifs from the CELF, RBFOX, and MBNL families are among the most frequently enriched across tumor types, as well as in luminal breast tumors (Fig. 4A) (Supplemental Figs. 14–18). Downregulated RBP genes in inclusion events and upregulated ones in skipping events show more frequently enriched motifs in upstream and exonic regions, consistent with a positional effect (Fig. 4B). On the other hand, downregulated RBP genes in skipping events and upregulated ones in inclusion events show enriched motifs most frequently on exons, suggesting that RBPs more often enhance the inclusion of bound exons.

**Figure 4.**
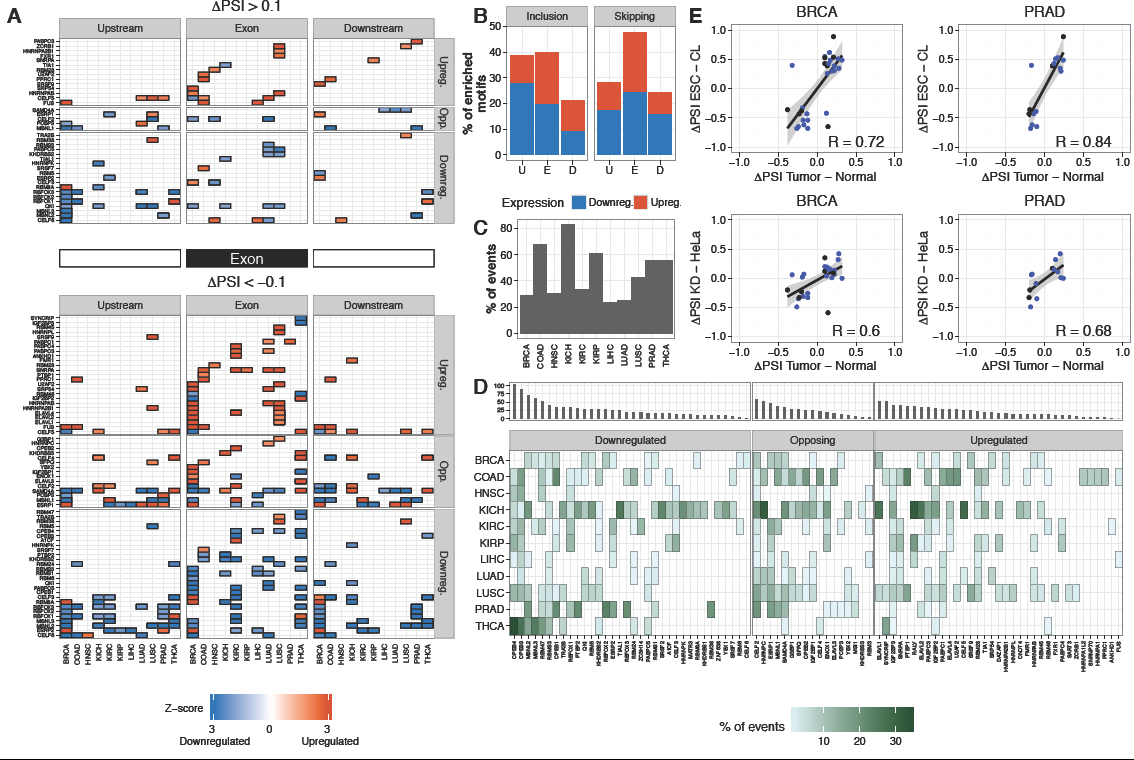
Enriched RNA binding motifs in differentially spliced events. **(A)** Enriched RNA binding motifs in differentially spliced skipping exon events in each tumor type, separated by inclusion (upper panels) or skipping (lower panels) events, and by upstream (left), exonic (middle) or downstream (right) regions. Only enriched motifs for splicing factors that are differentially expressed are indicated in each tumor type. RBPs (y axis) are grouped according to Fig. 1A. Gene up- and downregulation is indicated in red and blue, respectively. The color intensity indicates the Z-score of the motif enrichment. Similar plots for the other event types are given in the Supplemental Figs. S14-S18. **(B)** Proportion of enriched motifs in inclusion (ΔPSI > 0.1) (left panel) and skipping (ΔPSI < 0.1) (right panel) events, in each of the event regions (x-axis): upstream (U), exon (E) and downstream (D). Proportions are separated according to whether the RBP gene is up (red) or down (blue) regulated. **(C)** Total proportion (y-axis) of differentially spliced events in each tumor type (x-axis) that are predicted as targets of one or more differentially expressed RBPs with significance z-score > 1.96. **(D)** Proportion of differentially spliced events (marked in green) that are predicted as targets of each RBP (x axis) in each tumor type (y axis), with significance Z-score > 1.96. RBPs are grouped according to Fig. 1A. Upper panels indicate the number of unique cancer drivers with differentially spliced events predicted as targets of each RBP across all tumors. RBPs are ordered according to this value within each group. **(E)** Correlation (Pearson R) of ΔPSI values (x axes) in breast tumors (BRCA) and prostate tumors (PRAD) with the ΔPSI of events (y axes) from the knockdown of *MBNL1* and *MBNL2* in HeLa cells (upper panels) and from the comparison of stem cells (ESCs) with differentiated cells (CL) (lower panels). Events with a predicted MBNL binding motif are indicated in blue.

To define candidate target events for differentially expressed RBPs, we selected differentially spliced events whose PSI correlate with the RBP expression (|R| > 0.5, Spearman) and contain the corresponding RNA binding motif. We could assign between 20 and 80% of the differentially spliced events to at least one RBP (Fig. 4C) (Supplemental Tables S16 and S17). We ranked the RBPs in each expression group according to the total number of cancer drivers with differentially spliced events across all tumor types (Fig. 4D) (Supplemental Fig. S19). The results of this analysis emphasize the relevance of *CELF2, ESRP1, ELAVL1, RBFOX2, PTBP1, QKI, TRA2B* and *RBM47* in relation to alterations of splicing in cancer; and highlights novel prominent factors, including *HNRNPC, HNRNPQ (SYNCRIP)*, and genes from the *CPEB* and *MBNL* families. We confirmed the role *of MBNL1, QKI, RBFOX2, PTBP1, RBM47* and *ESRP1* in some of these tumors by comparing the ΔPSI values of the events with those obtained by single knockdown or overexpression experiments in cell lines (Fig. 4E, upper panels) (Supplemental Figs. S20-S22) (Supplemental Table S18). Additionally, comparing the tumor events with those differentially spliced between human embryonic stem cells (hESC) and differentiated cells or tissues (Han et al. 2013) yielded a high positive correlation of ΔPSI values in BRCA, PRAD, LIHC and breast luminal tumors; to a lesser extent in COAD and LUAD, and an anti-correlation in KIRC. This is in agreement with the *MBNL1* and *MBNL2* expression patterns in these tumors and with the majority of the correlating events containing the MBNL binding motif (Fig. 4E, lower panels) (Supplemental Fig. S23) (Supplemental Table S19). These results highlight the prominent role of RBP expression changes in the alterations of splicing in cancer and suggest new mechanisms of regulation.

### Network analysis uncovers overlapping RBP-mediated regulatory modules in cancer

To identify splicing regulatory networks relevant in cancer, we built clusters with the 162 SFs using the correlation between gene expression and event PSI values. We linked these clusters to differentially spliced genes in enriched cancer hallmarks (Methods). This analysis revealed modules of splicing regulation, with one or two genes as the main regulator of each hallmark across different tumor types. Myogenesis and EMT networks are enriched in almost all tumor types and controlled by similar factors (Figs. 5A and 5B) (Supplemental Figs. 24 and 25). Other relevant regulatory modules are the G2 checkpoint (G2M), which includes *NUMA1*, a gene involved in spindle formation (Zheng et al. 2010); and the WNT/Beta-catenin pathway, which includes *NUMB*, whose alternative splicing is linked to cell proliferation (Bechara et al. 2013). We predicted the splicing of these genes to be controlled by *MBNL1*, among other factors (Figs. 5C and 5D). Interestingly, angiogenesis, which is an enriched hallmark in COAD for splicing but not for gene expression, includes an event in *SERPINA5*, an inhibitor of serine proteases involved in homeostasis and thrombosis (Suzuki 2008), which we predict to be controlled by *RBM47, PTBP1* and *RBM28* (Fig. 5E). This analysis reveals new roles of RBPs and splicing in cancer-relevant processes.

**Figure 5.**
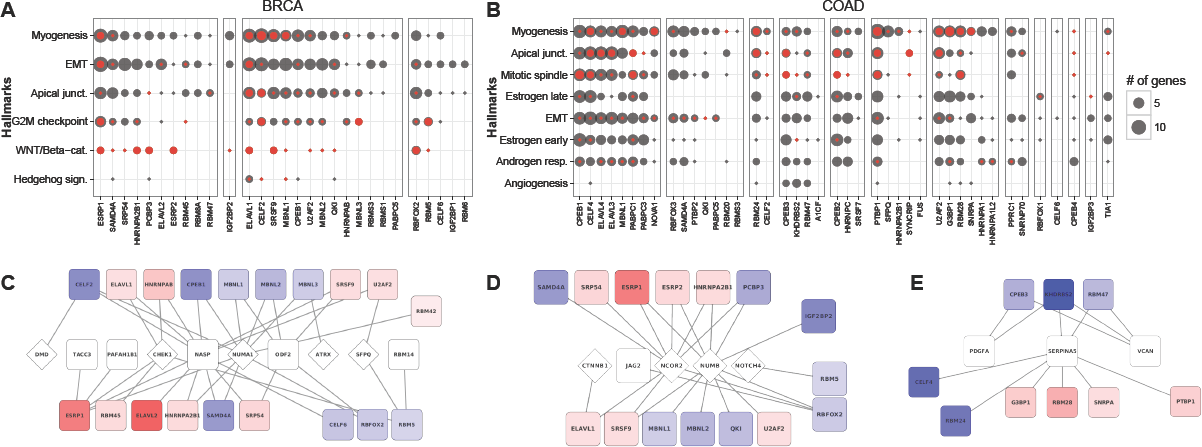
Networks of splicing regulation. Modules of alternative splicing regulation according to cancer hallmarks in breast **(A)** and colon **(B)** tumors. For each cluster of RBPs (x-axis) we indicate in gray the total number of genes targets linked to them in each hallmark (y-axis). Only enriched hallmarks are shown. We indicate in red the number of cancer drivers associated to each RBP, and in each cluster RBPs are ordered according to the total number of genes they are associated to. Representation of the regulatory modules for G2M checkpoint **(C)** and WNT/Beta-catenin **(D)** hallmarks in breast tumors, and for the angiogenesis hallmark **(E)** in colon tumors. RBPs are indicated as square boxes in red or blue depending of whether they are up-or downregulated. Target genes are presented as white diamonds for cancer drivers and white boxes for the rest. Connections indicate predicted splicing regulation by an RBP.

### MBNL1 contributes to cell proliferation through alternative splicing regulation of NUMA1

MBNL1 emerges as a potentially key regulator of splicing for multiple cancer drivers, especially in luminal breast tumors (Supplemental Fig. S26). In particular, MBNL1 is predicted to control an exon skipping event in *NUMA1*, whose PSI value correlates with *MBNL1* expression (Spearman R = 0.65/0.66 in luminal A/B) and contains the *MBNL1* binding motif (Fig. 6A) (Supplemental Fig. S27). The same event is more included in KIRC (ΔPSI = 0.11, corrected p-value = 3.11e-06), where *MBNL1* is significantly upregulated compared to normal tissues, providing further support for the dependence of *NUMA1* splicing on MBNL1. We detected MBNL1 protein in the breast epithelial cell line MCF10A, but not in the luminal-like MCF7 (Supplemental Fig. S28). *MBNL1* expression depletion in MCF10A cells with two different siRNAs induced skipping of exon 16 in *NUMA1*, recapitulating the pattern observed in the tumor samples (Fig. 6B, upper panel) and in MCF7 (Supplemental Fig. S28). We also tested the alternative splicing of *NUMB* exon 9, which we predicted to be dependent on *MBNL1* in BRCA luminal tumors. The depletion of *MBNL1* recapitulates *NUMB* splicing pattern in luminal samples (Fig. 6B, middle panel).

**Figure 6.**
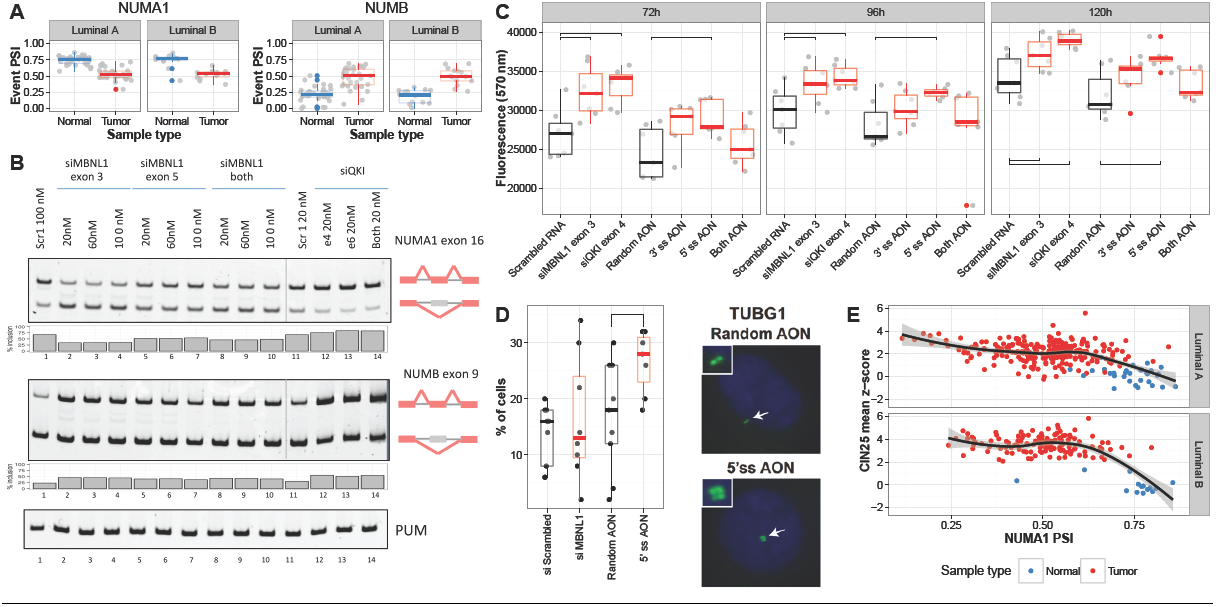
Regulation of NUMA1 alternative splicing by MBNL1 in breast luminal tumors. **(A)** PSI value distributions in tumor and paired normal sample for luminal A (LA) and luminal B (LB) breast tumors for the events in *NUMA1* (LA: ΔPSI = −0.22, p-value = 7.81e-07, LB: ΔPSI = −0.23, p-value = 0.037) and *NUMB* (LA: ΔPSI = 0.28 p-value = 0.0001, LB: ΔPSI = 0.28, p-value = 0.016). All p-values given are corrected for multiple testing. **(B)** Semi-quantitative RT-PCR isoform analysis upon knockdowns *of MBNL1* (lanes 2–10) and *QKI* (lanes 12–14) at different siRNA concentrations and the controls with scrambled siRNAs (lanes 1 and 11). The diagrams to the right indicate the position of the alternatively spliced exons. The bottom lanes correspond to the RT-PCR amplification of RNA from the PUM gene in the same samples, which are used as a control for RNA expression. **(C)** Resazurin-based assays of cell viability/proliferation. Measurements were performed in triplicate at 72, 96 and 120 hours. The plot shows measurements upon knockdowns *of MBNL1* (siMBNL1) and *QKI* (siQKI), upon transfection of AONs targeting the 3’ and 5’ splice-sites independently and both together, and the corresponding controls (scrambled siRNA and random AON). **(D)** Left panel: graph showing the results of the evaluation of centrosome amplification upon knockdown of *MBNL1* (siMBNL1) or upon transfection of AONs targeting 5’ splice-sites (5’ss AON), compared to the corresponding controls siScrambled (p=0,4271) and random AON (p=0,04356), respectively (one-sided Mann-Whitney test). Right panels: representative merged (TUBG1 and DAPI) images of immunofluorescence assays. **(E)** Correlation of *NUMA1* event PSI (x-axis) with the CIN25 signature of aneuploidy (y-axis) across the tumor (red) and normal (blue) samples for luminal A (upper panel) (R = −0.4 Spearman) and B (lower panel) (R = −0.33 Spearman).

For a comparison with *MBNL1*, we evaluated the role of *QKI*, which we also observed downregulated in BRCA luminal tumors and whose product was detected in MCF10A but not in MCF7 cells (Supplemental Fig. S28). Upon *QKI* expression depletion, *NUMA1* exon 16 inclusion changed by a small but reproducible amount in the direction opposite to that with *MBNL1* depletion (Fig. 6B, upper panel). Although we did not find a *QKI* motif on the *NUMA1* event, this is consistent with the negative correlation found with *QKI* expression (R = −0.11) in BRCA. Next, we also tested *NUMB* exon 9, which we predicted to be controlled by QKI in BRCA luminal tumors. *QKI* depletion induces exon 9 inclusion, recapitulating the pattern in luminal samples (Fig. 6B, middle panels). *NUMA1* alternative splicing is likely controlled by other factors (Fig 5C). For instance, *NUMA1* exon 16 contains an RBM42 binding motif and its PSI anti-correlates with *RBM42* expression in breast luminal tumors; hence *RBM42* potentially acts as a repressor of exon inclusion. Consistent with this, depletion of *RBM42* in MCF7 cells leads to increased inclusion of *NUMA1* exon 16 (Supplemental Fig. S28).

To measure whether the splicing change in *NUMA1* had any effect on cancer cell hallmarks, we used antisense oligonucleotides (AONs) targeting specifically the 5’ and 3’ splice-sites of *NUMA1* exon 16. These AONs promote exon skipping, recapitulating in the MCF10A cells the splicing pattern observed in BRCA luminal tumors, with the AON against the 5’ss being more efficient (Supplemental Fig. S29). We then measured the proliferation/viability of MCF10A cells transfected with the AONs targeting *NUMA1* exon 16, or with siRNAs against *MBNL1* and *QKI*. We observed a significant increase in cell proliferation/viability at 72, 96 and 120 hours upon depletion of *MBNL1* or *QKI* compared with controls (t-test p-value < 0.05) (Fig. 6C) (Supplemental. Table S20) and when transfecting cells with the AON against the 5’ splice site (t-test p-values < 0.05) (Fig. 6C). Using only the 3’ splice site or both AONs there was also an increase, albeit not statistically significant, possibly due to a smaller effect of the 3’AON on *NUMA1* splicing and the shared concentration of both AONs. Furthermore, although *QKI* has a mild effect on *NUMA1* exon 16 splicing, *QKI* downregulation has a much stronger effect on *NUMB* alternative splicing (Fig. 6B), which strongly promotes cell proliferation in a variety of cell types (Misquitta-Ali et al. 2011) (Bechara et al. 2013).

We further studied the possible effects of the alternative splicing of *NUMA1* exon 16 on centrosome amplification, a hallmark of breast carcinogenesis. Using the AON against the 5’ splice-site, there is a significant increase in number of cells with centrosome amplification compared with controls in MCF10A cells (Fig. 6D) (Supplemental Table S21). In contrast, the siRNA against *MBNL1* did not yield a significant difference, possibly due to the superposition of indirect effects. We also observed an inverse correlation between a signature for chromosome instability (Carter et al. 2006) with the *NUMA1* event PSI in luminal tumors (Fig. 6E), which was absent in other tumor types (Supplemental Fig. S30), providing further support to a relation between *NUMA1* alternative splicing and the fidelity of centrosome formation. The exon skipping described in *NUMA1* is the only coding difference between the tumor and the normal isoforms. While it is unclear whether this 14 amino acid change alone could explain the observed effects, e.g. we could not detect any protein domain or disordered region (Dosztányi et al. 2005), using GPS (Xue et al. 2011) we predicted loss of a high scoring threonine phosphorylation site (FDR **≤** 2%) upon exon skipping, suggesting a possible mechanism for the differential activities of the two isoforms (Supplemental Fig. S30) (Supplemental Table S22).

## Discussion

Our study reveals that alterations in genes encoding RNA binding proteins (RBPs) are pervasive in cancer, characterize the different tumor types and can explain many of the alternative splicing changes observed in tumors. Many of the RBP expression changes appear related to large copy number alterations. In contrast, mutations RBPs are particularly low in frequency compared to mutation rates in other genes. We measured the potential relevance of RBP alterations by looking at the associated splicing changes. Only a few RBPs beyond the previously described cases show mutations associated with genome-wide effects on alternative splicing. One of the novel cases is *HNRNPL*, for which we predict that frequent indels in an RNA binding domain would affect the splicing of *CASP8*, a gene involved in programmed cell death (Yang et al. 2008). As *HNRNPL* also controls *CASP9* alternative splicing (Goehe et al. 2010), this may suggest a relevant role *of HNRNPL* in apoptosis. On the other hand, the expression changes in RBP genes appear as major contributors of the alternative splicing changes observed in tumors, since the number of events affected by RBP mutations is lower than those changing between tumor and normal samples. The splicing changes detected are predicted to have an impact on many cancer hallmarks independently of changes in gene expression. This agrees with previous reports on the impact of splicing on cancer hallmarks (Oltean and Bates 2013) and highlights the relevance of alternative splicing as a complementary molecular mechanism to explain tumor development. An important implication of our study for prognostic and clinical studies is that the definition of functional impact of somatic genetic and epigenetic alterations should be expanded to include changes in the alternative splicing of the genes in cancer relevant pathways.

We identified many RBPs whose expression alteration has some potential relevance in cancer. For instance, *TRA2B* is frequently amplified and upregulated in LUSC, and its binding motif enriched in differentially spliced events, including an SE event in the DNA damage response gene *CHEK1*, reported recently to be controlled by Tra2 proteins (Best et al. 2014). Despite the similar genetic alterations between squamous tumors (Hoadley et al. 2014), only LUSC shows overexpression of *TRA2B*, which could explain some of the differences in splicing patterns between these two tumor types. We also found that genes from the MBNL family are frequently downregulated in tumors and their binding motif is enriched in differentially spliced events, especially in breast and prostate tumors. Additionally, splicing changes in the same tumor types recapitulate the splicing patterns of undifferentiated cells, in agreement with recent studies describing *MBNL1* and *MBNL2* as regulators of a stem cell related splicing program (Han et al. 2013; Venables et al. 2013). *MBNL1* potentially controls multiple genes that participate in cancer-related pathways, including the mitotic gene *NUMA1*. Although the *NUMA1* locus was related before to breast cancer risk (Kammerer et al. 2005), a clear mechanism explaining its relevance in cancer is still lacking. We described that *NUMA1* alternative splicing leads to higher proliferation and increased centrosome amplification in normal cells. *NUMA1* produces a protein component of the nuclear matrix, which is dependent on threonine-phosphorylation to regulate the orientation of mitotic spindles and ensure symmetric cell division (Kotak et al. 2013). *NUMA1* alternative splicing removes a potential threonine phosphorylation site; hence, one attractive possibility is that this splicing change affects NUMA1 phosphorylation, thereby impairing correct spindle positioning, leading to increased genome instability. Besides *MBNL1*, our analyses provided evidence for the control of *NUMA1* splicing by RBM42, in agreement with its proposed role in cell cycle (Suvorova et al. 2013) and pointing to an involvement of this factor in alternative splicing and cancer worth investigating further.

The origin of the expression changes in RBPs remains to be described. In particular, those cases that cannot be explained by DNA alterations in the gene locus may originate from alterations in the pathways that control the regulation of RBPs. It was recently postulated that RBPs involved in development and differentiation controlled by common enhancers could act as master regulators (Jangi and Sharp 2014). These included *MBNL1, RBFOX2, RBM24, RBM38, RBM20, RBFOX1, ZNF638*, and *RBMS3*, which we found frequently downregulated in tumors, with enriched motifs in differentially spliced events and potential targets in multiple cancer drivers. This suggests a general mechanisms in tumors by which the deactivation of one or more RBPs through the alteration of a common enhancer would lead to the reversal of multiple RNA splicing patterns to trigger or sustain an undifferentiated phenotype. Taken together, our results provide a rich resource of information about novel networks of RBPs that trigger common and specific alternative splicing changes in several solid tumors. These represent a set of candidate alternative splicing changes that may be relevant to understand the molecular basis of, and potentially reverse, the oncogenic properties of tumor cells.

## Methods

### Datasets

Datasets were downloaded from the TCGA data portal (https://tcga-data.nci.nih.gov/tcga/) (Supplemental Table S1). Details can be found in the Supplemental Methods. The 1348 genes coding for RNA binding proteins (RBPs) analyzed includes those with high confidence for RNA binding (Baltz et al. 2012; Castello et al. 2012; Kwon et al. 2013) and those annotated as RNA-binding in Ensembl (Cunningham et al. 2014) (Supplemental Table S2). From this set, a subset of 162 known and potential auxiliary splicing factors (SFs) was selected.

### Differential expression

Quantile normalization and voom (Law et al. 2014) transformation was performed on gene read counts. Differential expression was analyzed using the limma package (Smyth 2005) and p-values were corrected for multiple testing using the Benjamini-Hochberg method. Genes were considered differentially expressed if log2-fold change > 0.5 in absolute value and corrected p-value < 0.05. An expression Z-score per gene and per tumor sample was calculated using the quantile normalized and voom transformed read-counts.

### Mutation and Copy number variation analysis

The frequency of somatic mutations across all samples with available data was calculated per gene and tumor type. The association between expression regulation and mutations was measured using a Jaccard index. Mutual exclusion was measured using the number of samples having an RBP mutation and no expression change and the number of samples having expression change but no RBP mutation. For samples with CNV data available, the overlap of each RBP with the annotated CNVs was calculated, requiring a CNV score > log_2_(3) or < log_2_(1) for gain or loss, respectively. We required that the full locus of the RBP fall within a copy number region and defined focal CNVs to be those smaller than 5Mb. Using a Jaccard index, an association was calculated between the expression up- or downregulation and CNV gain or loss, respectively. More details are provided in the Supplemental Methods.

### Alternative splicing events

A total of 30820 alternative splicing events were calculated from the gene annotation using SUPPA (Alamancos et al. 2015): 16232 exon skipping (SE) events, 4978 alternative 5’ splice-site (A5) events, 6336 alternative 3’ splice-site (A3) events, 1478 mutually exclusive exon (ME) events, and 1787 retained intron (RI) events. Differentially spliced events were obtained by comparing the event percent spliced-in (PSI) value distributions between normal and tumor samples using a Wilcoxon signed rank test, removing samples with missing PSI values, using at least 10 paired samples, and correcting for multiple testing with the Benjamini-Hochberg method. Events were considered differentially spliced if the difference between the tumor and normal median PSIs > 0.1 in absolute value and a corrected p-value < 0.05. Events associated with mutations in RBPs were calculated in a similar way separating samples according to whether they had or not mutations in an RBP. Tests were performed using protein-affecting mutations. Only RBPs with mutations in at least 10 samples were tested. An enrichment Z-score was calculated per RBP and tumor type by comparing the number of events changing significantly with the median value obtained using all RBPs tested. Tumor type specific alternative splicing events were calculated by comparing their PSI values pairs of tumor types using an information theoretic measure. More details of the analyses are given in the Supplemental Methods.

### Gene sets

Annotations for 50 cancer hallmarks were obtained from the Molecular Signatures Database v4.0 (Liberzon et al. 2015) and a Fisher exact test was performed per hallmark using genes with annotated events and genes with differentially spliced events in each tumor type. The lists of 82 genes whose alternative splicing was linked before to cancer (Supplemental Table S5) and of 889 cancer drivers based on mutations and CNVs (34 in common with previous set) (Supplemental Table S6) were collected from the literature (more details in the Supplemental Methods).

### RNA binding motif enrichment

Details on how motifs from RNAcompete (Ray et al. 2013) were assigned to the RBP genes are given in the Supplemental Methods. The tool fimo (Bailey and Elkan 1994) was used to scan the motifs in the event regions using p-value < 0.001. Motif enrichment analysis was performed by comparing the frequency of regions in differentially spliced events with a motif with 100 random subsamples of the same size from equivalent regions in non-differentially spliced events controlling for similar G+C content. An enrichment Z-score per motif, region and direction of change (ΔPSI > 0.1, ΔPSI < −0.1) was calculated from the observed frequency and the 100 random control sets. A differentially spliced event was considered to be a potential target of a differentially expressed RBP if the correlation between the event PSI value and the gene expression robust Z-score was |R| > 0.5 (Spearman) and the event contained the RBP binding motif. To assess significance, the same number of differentially spliced events in a tumor type was randomly selected from all events 100 times, and events associated to the RBPs calculated each time as described previously. A Z-score was calculated from the mean and standard deviation. Cases with Z-score > 1.96 were considered significant.

### Networks of RBPs and events

Networks of RBPs and events were built using the correlations between RBPs through events. For each pair of RBP a correlation was calculated using the Spearman correlation values with all differentially spliced events in the same tumor type. RBP clusters were built by calculating an inverse covariance matrix of these correlations using the glasso algorithm (Friedman et al. 2008) and then searching for dense, highly connected sub-graphs with a greedy algorithm (Clauset et al. 2004). Events were associated to a network if they had |R| > 0.8 (Spearman) or |R| > 0.5 plus motif for any of the RBPs in an RBP cluster.

### Experimental procedures

Details on the cell cultures, siRNA transfections, Western blot analyses and semi quantitative RT-PCR experiments are given in the Supplemental Methods.

### Antisense oligonucleotides treatment

2’-O-Methyl RNA oligonucleotides were designed with full phosphorothioate linkage, antisense to the 5’ or 3’ splice sites of NUMA1 alternative exon 16 (hg19 coordinates chr11:71723447–71723488) optimizing GC content to 45–60 %. Custom modified and HPLC purified RNA oligos were ordered in a 0.2µM scale from SIGMA-ALDRICH. NUMA_1_ex16_5’ss: 5’- ggcauuacCUGCUUAGUUUGC-3’ NUMA1_ex16_3’ss: 5’- CCUCUAGCUGCUCCACcugu-3’ RANDOM 2’-O-Methyl RNA oligo: 5’-GCAAUGGCGUCAAGUGUGUCG-3’ Antisense RNA oligos were transfected in triplicate at 20nM final concentration using 2µl Lipofectamine RNAiMax (Life Technologies, 13778150) per 1ml of total volume of transfection in OPTIMEM (Life Technologies, 13778150). After five hours of treatment, media was replaced by DMEM-F12 containing 10% FBS and Pen-Strep.

### Cell proliferation/viability assay

2500 MCF10A cells/well were seeded the night before treatment in 96-well plates (NUNC, 167008) in 100µl complete DMEM-F12 medium. Wells with none, half or double amount of cells were also seeded for fluorescence calibration. Cells were transfected with siRNA or AON oligos as described. Resazurin (SIGMA, R7017) treatment was performed 72, 96 and 120 hours after transfection, in 7 replicates and incubated for 4 hours in a 37ºC incubator. Fluorescence was measured after 4 hours of incubation, using a TECAN infinite m200 device with 530 nm excitation wavelength, 590 nm emission wavelength, 30 nm emission bandwidth, and set to optimal gain. The medium was replaced by complete DMEM-F12 after measurements.

### Centrosome count and aneuploidy signature

The number of centrosomes was determined by immunofluorescence assays using an anti-γ-tubulin (TUBG1) antibody (clone GTU-88, Sigma-Aldrich; dilution 1:1,000). The expected immunostaining pattern of this centrosomal marker in normal cells is one or two foci proximal to the nucleus. The cells were fixed in cold methanol for 10 minutes and washed in phosphate-buffered saline. The secondary antibody was Alexa Fluor 488 (Molecular Probes, Life Technologies) and the cells were mounted using VECTASHIELD^®^ with DAPI. The results correspond to at least five independent fields and > 200 cells analyzed. The significance of the results was assessed using the one-sided Mann-Whitney test (Supplemental Table S21). The chromosome instability signature CIN25 (Carter et al. 2006) was used by calculating the mean value of the normalized expression robust Z-score values for the 25 genes from the signature in each sample.

## Acknowledgements

We are thankful to P. Papasaikas, B. Blencowe, M. Irimia and Q. Morris for comments and discussions. ES, BS, AP and EE were supported by the MINECO and FEDER (BIO2014-52566-R), Consolider RNAREG (CSD2009-00080), by AGAUR (SGR2014-1121) and by the Sandra Ibarra Foundation for Cancer (FSI2013). JV and BM were supported by Fundación Botín, by Banco de Santander through its Santander Universities Global Division and by Consolider RNAREG (CSD2009-00080), MINECO and AGAUR. FM and MAP were supported by AECC (Hereditary Cancer), AGAUR (SGR2014-364), the ISCIII, the MINECO and FEDER (PIE13/00022-ONCOPROFILE, PI15/00854 and RTICC RD12/0036/0008).

